# Imaging tissues and cells beyond the diffraction limit with structured illumination microscopy and Bayesian image reconstruction

**DOI:** 10.1101/426296

**Authors:** Jakub Pospíšil, Tomáš Lukeš, Justin Bendesky, Karel Fliegel, Kathrin Spendier, Guy M. Hagen

## Abstract

**Background:** Structured illumination microscopy (SIM) is a family of methods in optical fluorescence microscopy that can achieve both optical sectioning and super-resolution effects. SIM is a valuable method for high resolution imaging of fixed cells or tissues labeled with conventional fluorophores, as well as for imaging the dynamics of live cells expressing fluorescent protein constructs. In SIM, one acquires a set of images with shifting illumination patterns. This set of images is subsequently treated with image analysis algorithms to produce an image with reduced out-of-focus light (optical sectioning) and/or with improved resolution (super-resolution).

**Findings:** Five complete and freely available SIM datasets are presented including raw and analyzed data. We report methods for image acquisition and analysis using open source software along with examples of the resulting images when processed with different methods. We processed the data using established optical sectioning SIM and super-resolution SIM methods, and with newer Bayesian restoration approaches which we are developing.

**Conclusion:** Various methods for SIM data acquisition and processing are actively being developed, but complete raw data from SIM experiments is not typically published. Publicly available, high quality raw data with examples of processed results will aid researchers when developing new methods in SIM. Biologists will also find interest in the high-resolution images of animal tissues and cells we acquired. All of the data was processed with SIMToolbox, an open source and freely available software solution for SIM.

## Data description

### Context

Several methods are now available which are able to extend the resolution of fluorescence microscopy beyond the diffraction limit. These methods include photoactivated localization microscopy [1,2] (PALM, FPALM), stochastic optical reconstruction microscopy [3,4] (STORM, dSTORM), super-resolution optical fluctuation imaging [5,6] (SOFI), stimulated emission depletion microscopy [7] (STED), and structured illumination microscopy [8,9] (SIM).

Of these various methods, SIM is usually regarded as the most useful for imaging live cells, and this method has rapidly gained in popularity. Depending on the optical setup and data processing method used, SIM can achieve optical sectioning (OS-SIM) [10], an effect which greatly reduces out-of-focus light similar to laser scanning confocal fluorescence microscopy. SIM can also be used for imaging beyond the diffraction limit in fluorescence microscopy. Super-resolution SIM (SR-SIM) [8,9], in its most common implementation [11], uses laser illumination to create a high frequency interference fringe pattern (close to or at the resolution limit of the microscope) to illuminate the sample. In such an experiment, image information with details beyond the limit of spatial frequencies accepted by the microscope is aliased into the acquired images. By acquiring multiple images with shifting illumination patterns, a high-resolution image can be reconstructed [8,9]. Two-dimensional SR-SIM enables a twofold resolution improvement in the lateral dimension [8,9,12,13]. If a three-dimensional illumination pattern is used, a twofold resolution improvement can also be realized in the axial direction [11,14,15]. SIM is perhaps the most attractive super-resolution method for imaging live cells because it does not require high illumination powers, can work with most dyes and fluorescent proteins, uses efficient widefield (WF) detection, and can achieve high imaging rates. SIM has been demonstrated in several applications, including 2D [12,13], and 3D imaging [14,16].

As interest in super-resolution imaging has increased, several alternative approaches for SIM have been introduced which use various kinds of patterned illumination [17–21]. For example, in multifocal structured illumination microscopy (MSIM) [17], a 2D array of focused laser spots is scanned across a sample, and subsequent image processing is used to achieve an image with improved resolution. Structured illumination methods have also been combined with light sheet excitation, a method ideal for imaging live cells [22–26].

In addition to new illumination schemes, alternative data processing methods have also been introduced [27–33]. For example, Orieux et al. suggested a 2D method for SIM reconstruction based on Bayesian estimation [28], and our group showed that Bayesian reconstruction methods in SIM have several potential advantages and can achieve a performance comparable to traditional SIM methods [29]. To allow 3D imaging, our group subsequently introduced maximum *a posteriori* probability SIM (MAP-SIM [30]), a method based on reconstruction of the SIM data using a Bayesian framework. Image restoration approaches are useful when working with low signal levels in SIM [34], and have been recently reviewed [35].

We present complete raw and analyzed SIM data from several different situations in cell biology studies in which we imaged both live and fixed mammalian cells as well as fixed tissues. We used an alternative approach for SIM illumination which has been previously described [30,36,37]. Our system uses either light emitting diode (LED) or laser illumination, and a fast ferroelectric liquid crystal-on-silicon (FLCOS) microdisplay (also known as a spatial light modulator (SLM)) for SIM pattern definition. SLMs have seen use in SIM and related applications when high speed imaging and flexibility in controlling the spatial and temporal properties of the illumination are priorities [12–14,16,25,37–43]. To analyze the data we used OS-SIM, SR-SIM, and MAP-SIM methods. All of the raw and analyzed data are available on GigaDB, and the analysis software (SIMToolbox) is open-source and freely available [36].

## Methods

### Cell lines and reagents

All cell lines used were maintained in DMEM supplemented with 10% FCS, 100 U/ml penicillin, 100 U/ml streptomycin, and L-glutamate (Invitrogen) at 37 °C and 100% humidity. Cell lines we used for this study included U2-OS (human bone sarcoma), A431 (human skin carcinoma), and Hep-G2 (human liver carcinoma).

### Preparation of samples for imaging

(SIM data 1, Fig. 4) U2-OS cells expressing lysosome-associated membrane protein 1 labeled with green fluorescent protein (LAMP1-GFP) were grown in petri dishes with coverslip bottoms (MatTek) for 24 hours, then imaged them in full medium at room temperature. In this experiment, we used microscopy system 1 (Olympus IX71, Table 2).

(SIM data 2, Fig. 5) A431 cells were grown on #1.5H coverslips (Marienfeld) for 48 hours in normal medium. We washed the cells once with phosphate buffered saline (PBS), pH 7.4, and then treated the cells with 5 µM DiI-C_16_ (Molecular Probes) in PBS at room temperature for 5 minutes. This probe is a lipid modified with a fluorescent dye that inserts into the plasma membrane of live mammalian cells within a few minutes. We then washed the cells twice with PBS, then imaged them on the SIM system in fresh PBS at room temperature using a coverslip chamber (Biotech). In this experiment, we used microscopy system 3 (Leica DMi8, Table 2).

(SIM data 3, Fig. 6) A prepared slide was acquired (AmScope) containing sectioned rabbit testis stained with hematoxylin and eosin (H&E). In this experiment, we used microscopy system 3 (Leica DMi8, Table 2).

(SIM data 4, Fig. 7) Hep-G2 cells expressing Dendra2-histone 4 [44] were grown on #1.5H coverslips for 24 hours, then fixed for 15 minutes at room temperature with 4% paraformaldehyde. We then permeabilized the cells for 5 minutes at room temperature with 0.1% triton-X100, then washed the cells with PBS. We then labeled the actin cytoskeleton of the cells for 1 hour at room temperature with 5 nM Atto 565 phalloidin, followed by washing the cells with PBS. We finally mounted the coverslips on clean slides using mowiol 4-88 (Fluka). In this experiment, we used microscopy system 1 (Olympus IX71, Table 2).

(SIM data 5) A prepared slide was acquired (Molecular Probes) containing bovine pulmonary endothelial (BPAE) cells stained with Alexa Fluor 488 phalloidin (to label the actin cytoskeleton) and Mitotracker CMXRos (to label mitochondria). In this experiment, we used microscopy system 2 (Olympus IX83, Table 2).

Table 1 summarizes the imaging parameters used for the different samples.

**Table 1.** Imaging parameters for the SIM datasets.

| Data | sample | Label (structure) | Pixel size, nm | Illumination | Exposure time, ms | SIM experiment type | SIM pattern # of angles/phases | Microscope system used |
| --- | --- | --- | --- | --- | --- | --- | --- | --- |
| SIM data 1 (Fig. 4) | Live U2-OS cells | LAMP1-GFP (lysosomes and membrane) | 65 | LED 480 nm | 25 | 2D time lapse | 1/11 | 1 |
| SIM data 2 (Fig. 5) | Live A431 cells | Dil-C16 (membrane) | 65 | LED 530 nm | 100 | 3D | 4/24 | 3 |
| SIM data 3 (Fig. 6) | Fixed rabbit testis | Hematoxylin and eosin (structural strain) | 65 | LED 530 nm | 200 | 3D | 1/11 | 3 |
| SIM data 4 (Fig. 7) | Fixed Hep-G2 cells | Dendra2-H4 (nucleus)<br>Atto565-phalloidin (actin) | 65 | LED 480 nm<br>LED 530 nm | 500 | 3D | 4/24 | 1 |
| SIM data 5 (Fig. 8) | Fixed BPAE cells | AlexaFluor 488 phalloidin (actin)<br>Mitotracker CMXRos (mitochondria) | 65 | Lumencor spectra-X 470 nm<br>550 nm | 300 | 2D | 1/11 | 2 |

### Microscope setup and acquisition

We used three different home-built SIM setups based on the same general design as described previously [30,36,37] (Figure 1). The three SIM systems were based on Olympus IX71, Olympus IX83, and Leica DMi8 microscopes coupled with sCMOS cameras (Andor) under the control of IQ3 software (Andor). The parameters of the different microscope setups are shown in table 2.

**Figure 1:**
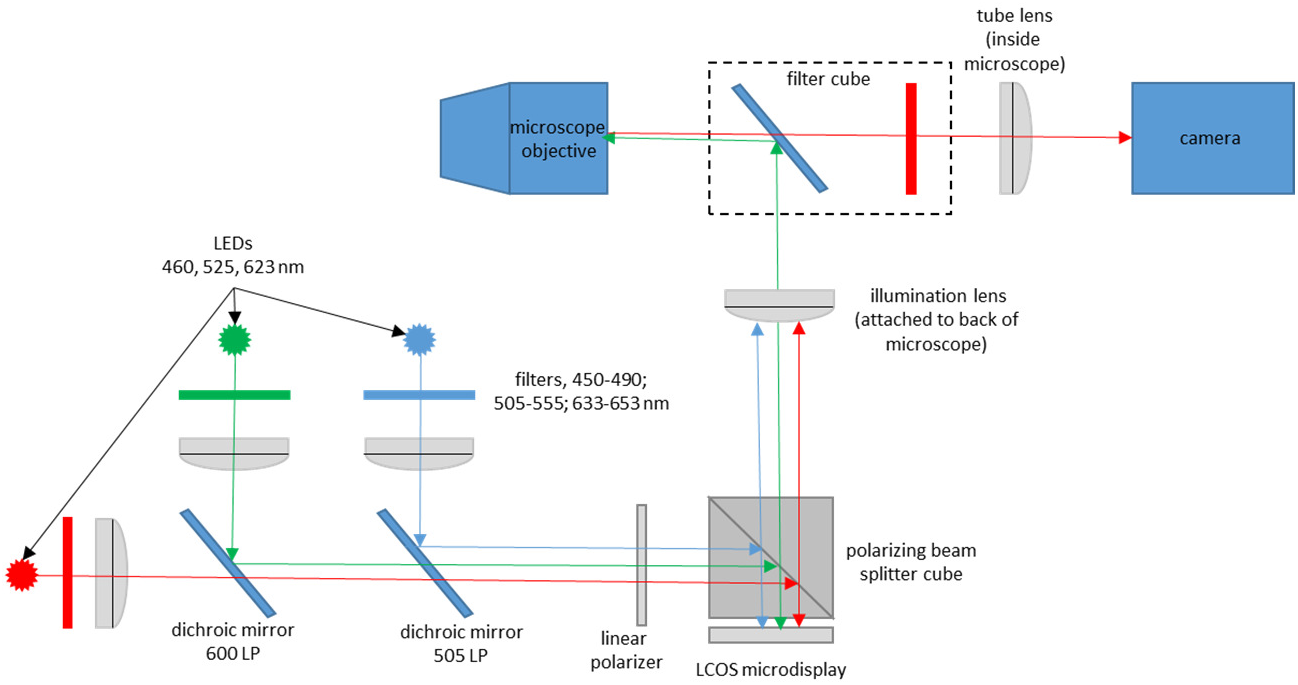
Structured illumination microscope setup, which we used with different microscope bodies and cameras. See text and table 2 for details.

**Table 2.** Parameters of the microscope systems.

| Setup | Microscope | Objective | sCMOS Camera | Illumination tube lens focal length and part number |
| --- | --- | --- | --- | --- |
| 1 | Olympus IX71 | 100×/1.4 UPLSAPO | Andor Neo 5.5 | 180 mm U-TLU |
| 2 | Olympus IX83 | 100×/1.3 UPLFLN | Andor Zyla 4.2+ | 180 mm SWTLU-C |
| 3 | Leica DMI8 | 100×/1.47 HCX PLAPO TIRF | Andor Zyla 4.2+ | 200 mm 11525408 |

In each microscope setup, the illumination patterns were produced by a high-speed ferroelectric liquid crystal on silicon (FLCOS) microdisplay (SXGA-3DM, Forth Dimension Displays, 13.6 µm pixel pitch). This particular FLCOS microdisplay has been used previously in SIM [14,16,25,29,30,36,37,45–48], and in other optical sectioning systems such as programmable array microscopy (PAM) [38,42,49]. The display was illuminated by a home-built, three channel LED system based on high power LEDs (PT-54 or PT-120 with DK-114N or DK-136M controller, Luminous Devices) with emission maxima at 460 nm, 525 nm, and 623 nm. The output of each LED was filtered with a band pass filter (Chroma), and the three wavelengths were combined with appropriate dichroic mirrors (Chroma). The light was then vertically polarized with a linear polarizer (Edmund Optics). We imaged the microdisplay into the microscope using an external tube lens (Table 2) and polarizing beam splitter cube (Thor Labs). With any of the setups and when using a 100× objective, single microdisplay pixels are imaged into the sample with a nominal size of 136 nm, thus as diffraction-limited spots. This is important for achieving the highest resolution results [37]. More details are available in the supplementary material of [36]. In one experiment (Fig. 8), we used a Spectra-X light source (Lumencor).

The microdisplay allows one to create any desired illumination pattern. In our experiments, the illumination masks consisted of line grids of different orientations (0°, 90°, 45° and 135°). The lines were one microdisplay pixel thick (diffraction limited in the sample when using a 100× objective) with a gap of “off” pixels in between. The illumination line grid was shifted by one pixel between each image acquisition to obtain a shifted illumination mask. The shift between each image was constant, and the sum of all illumination masks resulted in homogenous illumination. Our optical setup, in which an incoherently illuminated microdisplay is imaged into the sample with highly corrected microscope optics, results in much more stable SIM illumination parameters compared to conventional SIM in which the illumination pattern is created by laser interference. We use a unique spatial calibration method to determine, with very high accuracy, the position of the patterned illumination in the sample [37]. This is a spatial domain process and does not rely on fitting of data to a model except for the assumption that the imaging is linear and shift-invariant.

### Data processing methods

We processed all of the data presented here using SIMToolbox, an open source, user friendly, and freely available program which our group developed for processing SIM data [36]. SIMToolbox, sample data, and complete documentation are freely available (http://mmtg.fel.cvut.cz/SIMToolbox). SIMToolbox is capable of OS-SIM [10,37], SR-SIM [8,9], and MAP-SIM [30] methods. See the supplementary information for additional details about these methods.

### Resolution measurements - spatial domain method

Previously, we used microscopy setup 1 (Olympus IX71) to measure spatial resolution by averaging spatial measurements from fifty individual 100 nm fluorescent beads [30]. We used a 100×/1.40 NA oil immersion objective and 460 nm LED excitation (emission 500 - 550 nm). A 19 × 19 pixel region of interest (ROI) was selected around each bead in both the widefield and MAP-SIM images. The ROIs were then registered with sub-pixel accuracy using normalized cross-correlation. Each ROI was fit with a Gaussian function and the full width at half maximum (FWHM) was determined in the axial and lateral directions. Figure 2 shows the resulting averaged FWHM values and PSF cross-sections [30].

**Figure 2:**
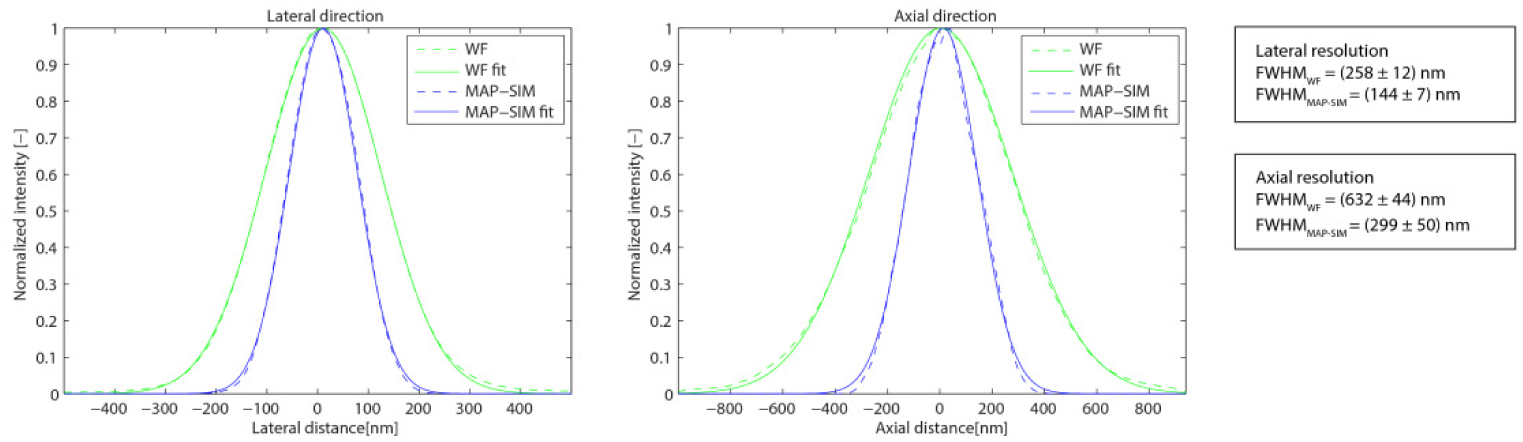
Measurements of the spatial resolution on a sample of fluorescent beads. Cross-sections of the PSF are obtained by averaging measurements over 50 beads along lateral and axial directions.

### Resolution measurements - frequency domain method

It is desirable to measure the actual resolution achieved in SIM images (or image sequences) of cells or tissues, but suitable structures are not always present in the images. We therefore developed a robust frequency domain method which can be used to measure resolution in any fluorescence microscopy image [50].

The power spectral density (PSD) describes the distribution of the power of a signal with respect to its frequency. The PSD of an image is the squared magnitude of its Fourier transform, and can be written as

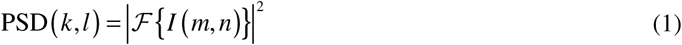

where 𝓕 represents the Fourier transform, *I(m*,*n)* is the image intensity, *m*,*n* indexes the rows and columns of the 2D image, respectively, and (*k*,*l)* are coordinates in the frequency domain. In polar coordinates, the circularly averaged PSD (PSD_ca_) in frequency space with frequency *q* and angle *θ* is given as

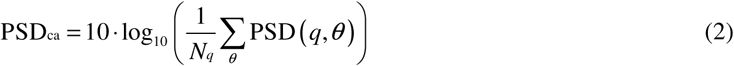

which averages PSD at spatial frequency *q*. *N_q_* is the number of pixels at a particular frequency *q*. The resolution limit in real space corresponds to the cut-off frequency in Fourier space. Assuming a noiseless case, the cut-off frequency will be equal to the spatial frequency at which PSD_ca_ drops to zero. In practice, PSD_ca_ contains non-zero values over the whole frequency range caused by noise. The signal to noise ratio (SNR) in Fourier space is generally very low close to the cut-off frequency, which makes precise detection of the cut-off frequency challenging. For this we use a spectral subtraction method [50]. Assuming additive noise, in the frequency domain we can write

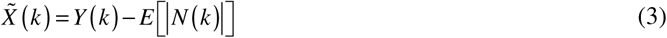

where
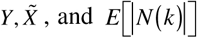
represent the noisy signal, the desired signal, and the noise spectrum estimate (expected noise spectrum), respectively. The amplitude noise spectrum |*N* (*k*)| is estimated from the parts of signal where only noise is present. If the spatial sampling is high enough to fulfill the Nyquist–Shannon criterion and oversamples the resolution limit of the reconstructed SIM image, spatial frequencies close to the half of the sampling frequency do not contain useful signal and can be used for noise estimation. We varied the frequency cut-off threshold over the range 〈0.95 *f*_max_; *f*_max_〉, estimated the level of noise for every threshold value, and obtained the mean and variance of the cut-off frequency (i.e. the resolution estimate). The *f*_max_ is given by
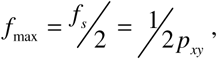
where *f_s_* and *p_xy_* are the sampling frequency and the backprojected pixel size, respectively.

Figure 3 shows the PSD_ca_ and corresponding resolution limit measured for the data shown in Fig. 5. Using our resolution estimation algorithm, we calculated a lateral spatial resolution of 294 nm for WF, and 141 nm for MAP-SIM. The measured resolution is in approximate agreement with our results measured on 100 nm fluorescent beads (Fig. 2).

**Figure 3:**
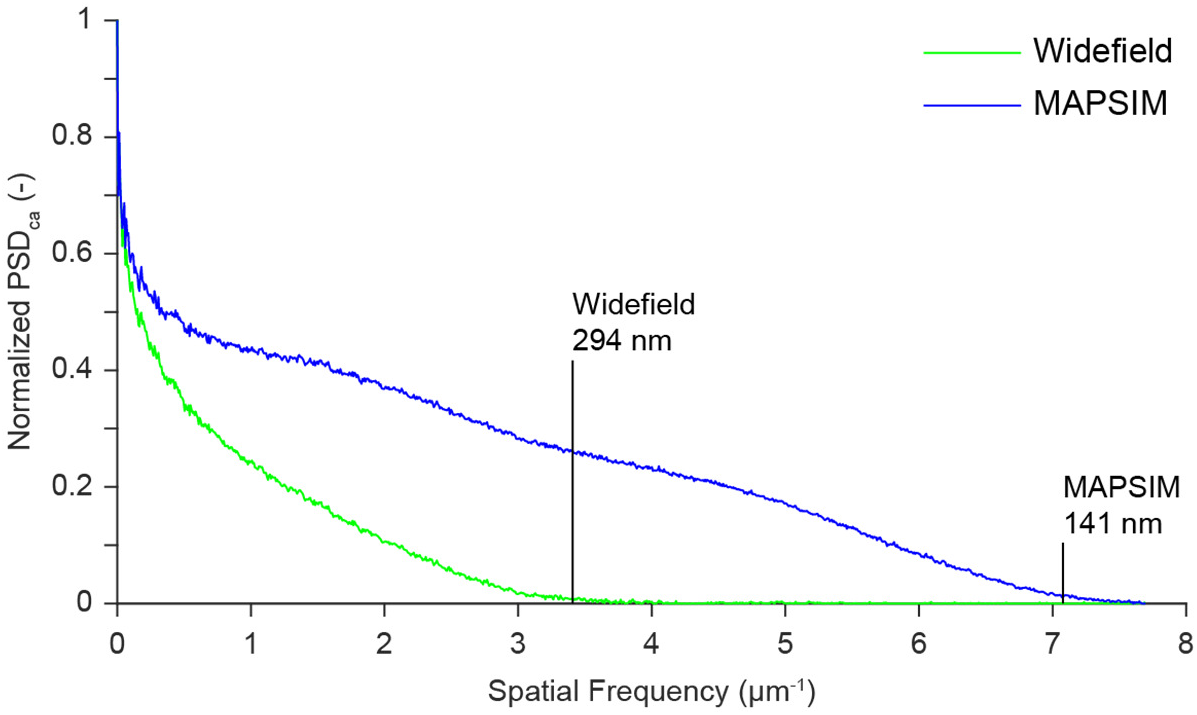
Resolution analysis and normalized power spectral density (PSD) measured on a selected image from the data in Fig. 5. The results indicate a circularly-averaged PSD lateral spatial resolution of 294 nm for WF, and 141 nm for MAP-SIM, in approximate agreement with the analysis in Fig. 4(d-f).

### Imaging live cells, fixed cells, and tissues with SIM

To demonstrate the utility of our approach in imaging live cells, we imaged U2-OS cells that had been transfected with GFP-tagged lysosomal associated membrane protein (LAMP1-GFP). LAMP1 is a highly glycosylated protein which is found on the surface of lysosomes and in the plasma membrane [51]. Fig. 4 shows widefield, OS-SIM, and MAP-SIM images of U2-OS cells expressing LAMP1-GFP, and the fast Fourier transform (FFT) of each image. The dotted circles in Fig. 4(d-f) show the approximate limit of resolution in each image. We found that, in addition to lysosomal expression, LAMP1-GFP is also present in high concentrations in the plasma membrane of U2-OS cells.

**Figure 4:**
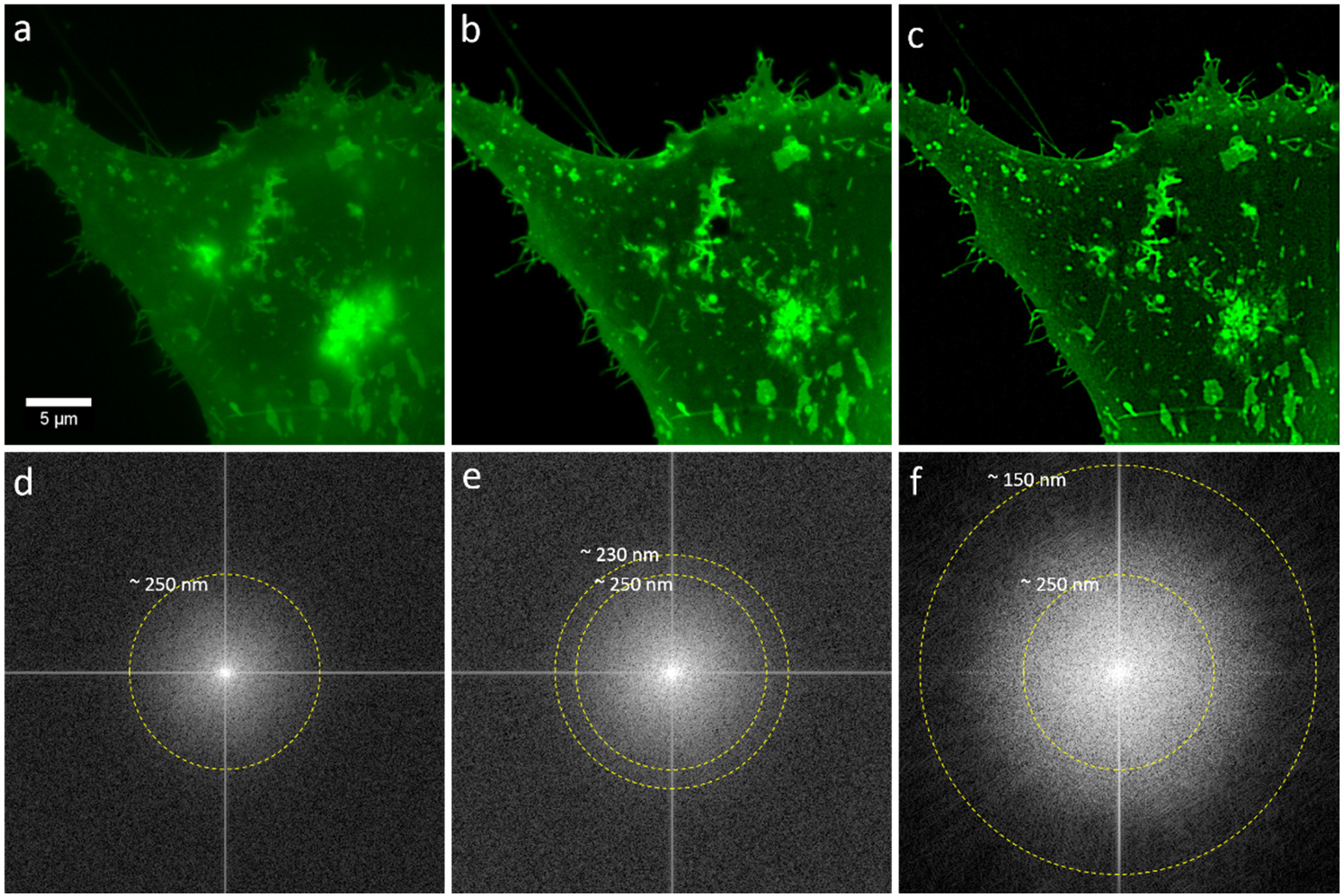
Imaging live cells beyond the difraction limit with MAP-SIM. U2-OS cells expressing LAMP1-GFP were imaged using the LCOS-based SIM system. Subsequent processing using OS-SIM or MAP-SIM methods. (a) WF, (b) OS-SIM, (c) MAP-SIM, (d) FFT of WF, (e) FFT of OS-SIM, (f) FFT of MAP-SIM. The images were individually scaled for presentation. The dotted cirular lines indicate the approximate resolution achieved in each image according to analysis of the FFT. The full image sequence is available at http://mmtg.fel.cvut.cz/mapsimlive_suppl/.

In this experiment, we acquired SIM image sequences with an exposure time of 25 ms, a raw imaging rate of 40 Hz. We used a SIM pattern with 11 phases (pattern period in the sample plane 1.5 µm) and a single angle (0° with respect to the camera), acquiring 3982 total frames, resulting in 472 processed frames (see table 1). The imaging rate of processed result frames was therefore 3.6 Hz. The full image sequence is available at http://mmtg.fel.cvut.cz/mapsimlive_suppl/. It is also available at Giga DB. We further analyzed this data as shown in the supplementary material (Figure S2-S3).

We next imaged live A431 cells which we labeled with the fluorescent lipid DiI-C16. In this experiment we acquired SIM image sequences with an exposure time of 100 ms, a raw imaging rate of 10 Hz. We used a SIM pattern with 24 total phases and four angles (see table 1). This data is shown in Figure 5.

**Figure 5:**
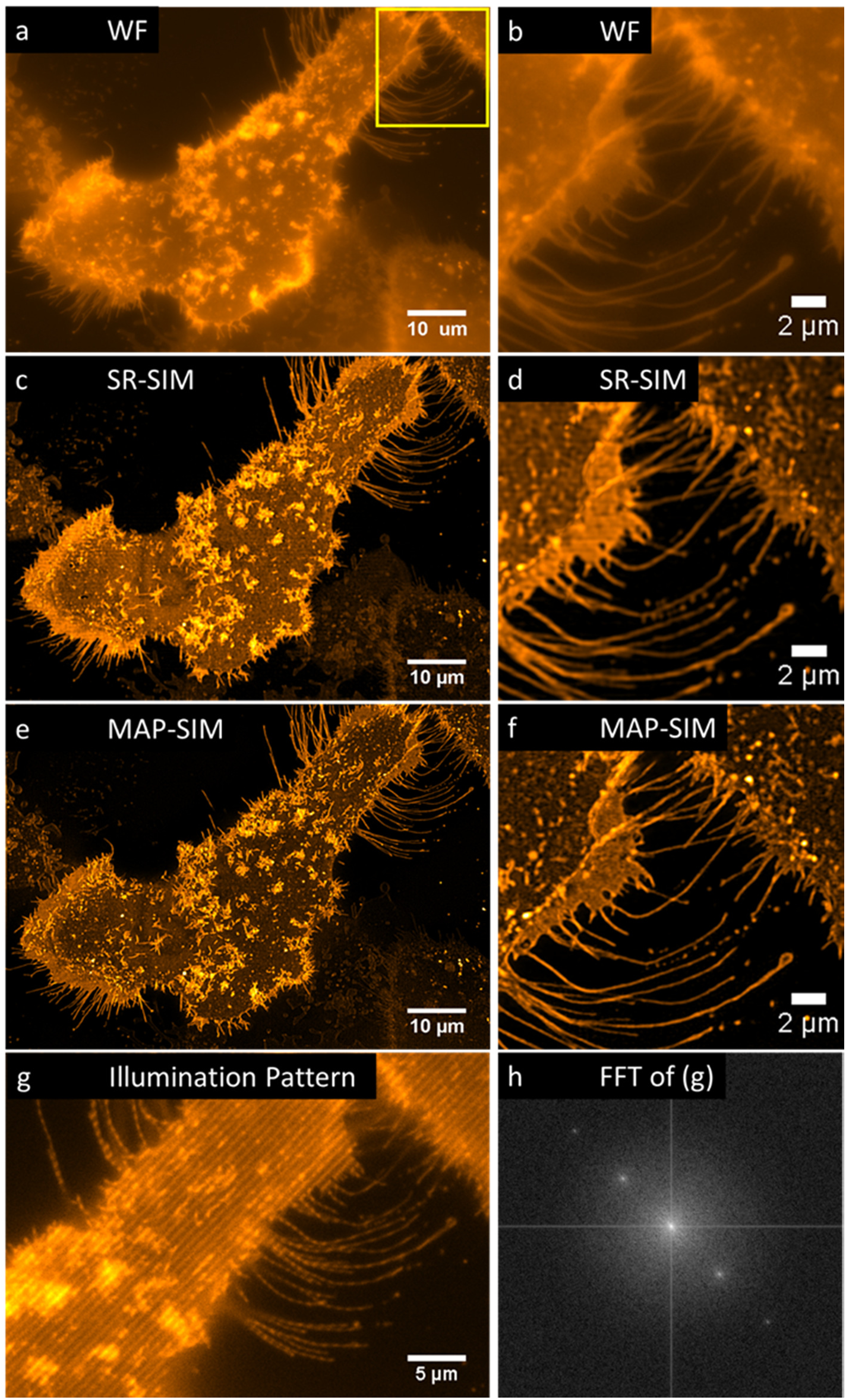
Imaging live cells beyond the difraction limit with SIM. A431 cells labeled with DiI-C16 were imaged using the LCOS-based SIM system. Subsequent processing using SR-SIM or MAP-SIM methods. (a) WF, (c) SR-SIM, (e) MAP-SIM. (b), (d), and (f) each show a zoom-in of the region indicated in (a). (g) shows the SIM illumination pattern in one of the four angles used. (h) shows a FFT of the image in (g). The images were individually scaled for visualization purposes. Each is a maximum intenstiy projection of 3 Z positions (spacing 400 nm (except for g and h which show a single Z-position).

Figure 6 shows SIM imaging of fixed tissues, in this case the seminiferous tubule of the rabbit stained with hematoxylin and eosin. Figure 7 shows SIM imaging of fixed HEPG2 cells expressing H4-Dendra, a nuclear marker. We also stained the cells with Atto 532-phalloidin to label the actin cytoskeleton. Figure 8 shows SIM imaging of fixed BPAE cells labeled with Alexa 488-phalloidin and mitotracker CMXRos to visualize the actin cytoskeleton and mitochondria, respectively.

**Figure 6:**
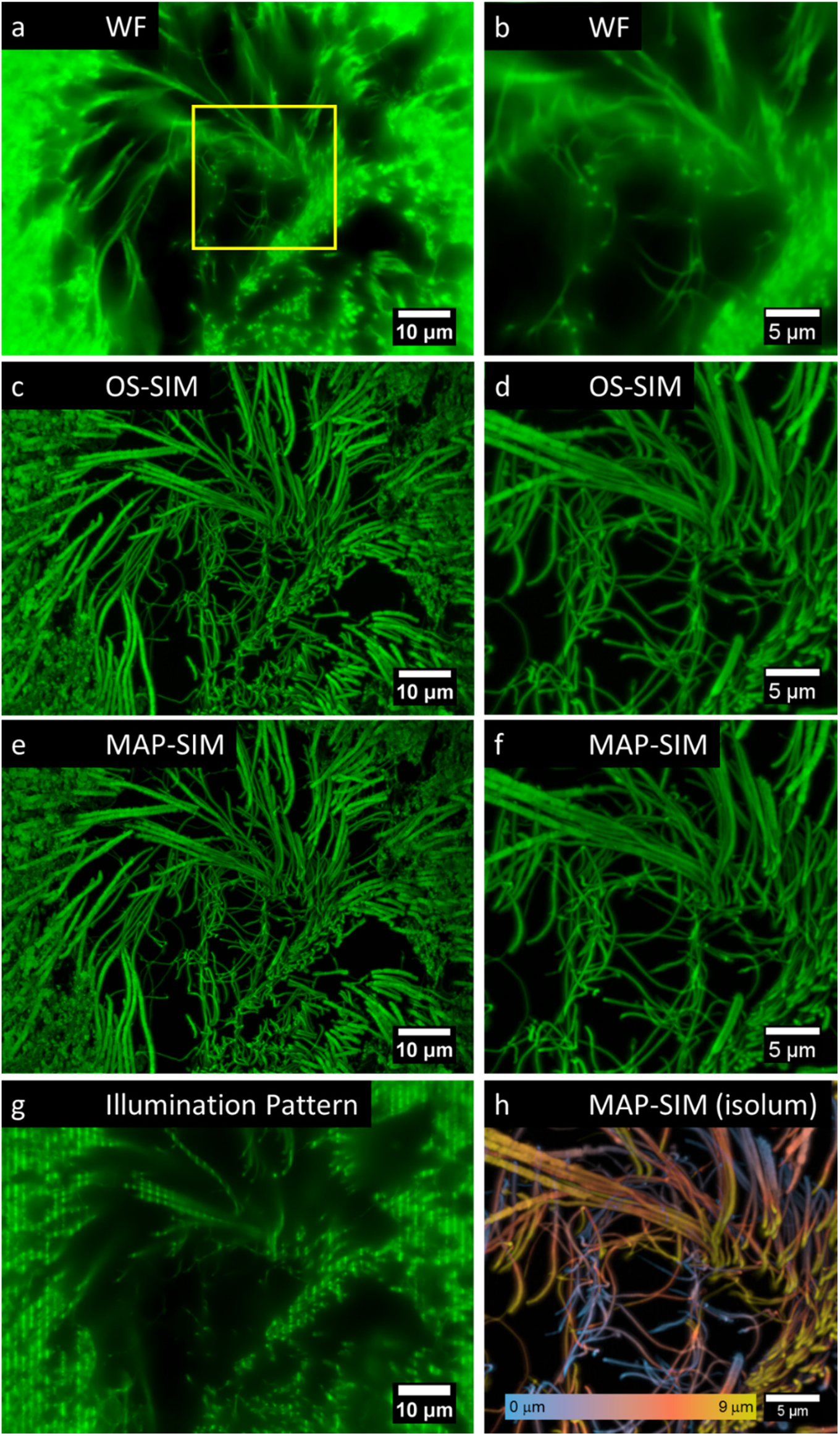
Imaging animal tissues using the LCOS-based SIM system and subsequent processing using OS-SIM or MAP-SIM methods. Seminiferous tubule of the rabbit stained with hematoxylin and eosin. (a) WF, (c) OS-SIM, (e) MAP-SIM. (b), (d), and (f) each show a zoom-in of the region indicated in (a). (g) shows the SIM illumination pattern in one of the four angles used. (h) MAP-SIM depth-coded using the lookup table isolum [52]. The images were individually scaled for visualization purposes. Each is a maximum intenstiy projection of 31 Z-positions (spacing 300 nm (except for (a, b, g) which show 1 Z-position).

**Figure 7:**
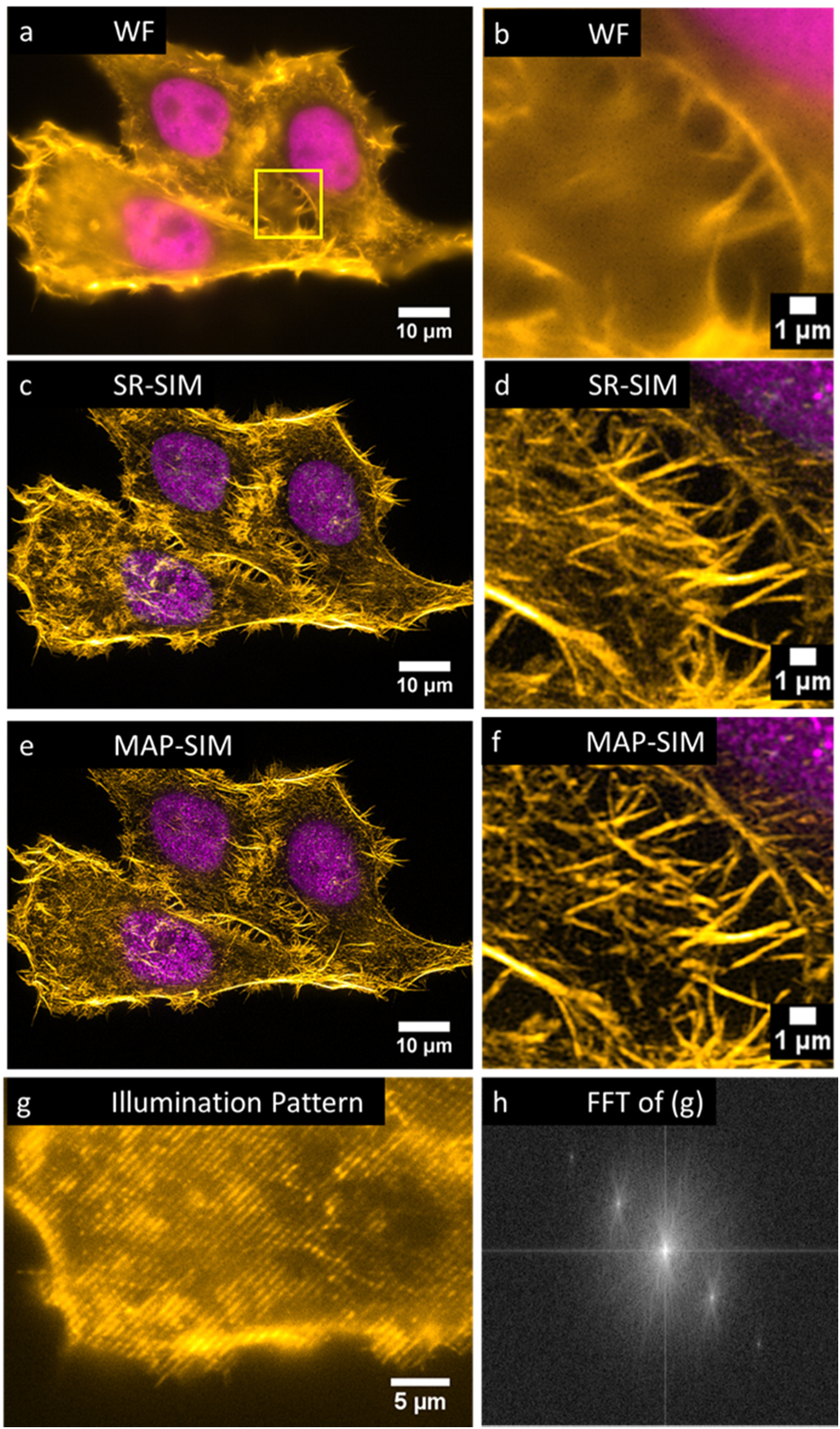
SIM imaging of fixed HEP-G2 cells expressing Dendra2-H4 (nucleus) and labeled with Atto-532 phalloidin. (a) WF, (c) SR-SIM, (e) MAP-SIM. (b), (d), and (f) each show a zoom-in of the region indicated in (a). (g) shows the SIM illumination pattern in one of the four angles used. (h) shows a FFT of the image in (g). The images were individually scaled for visualization purposes. Each is a maximum intenstiy projection of 22 Z-positions (spacing 200 nm (except for a, b, g and h which show 1 Z-position).

**Figure 8:**
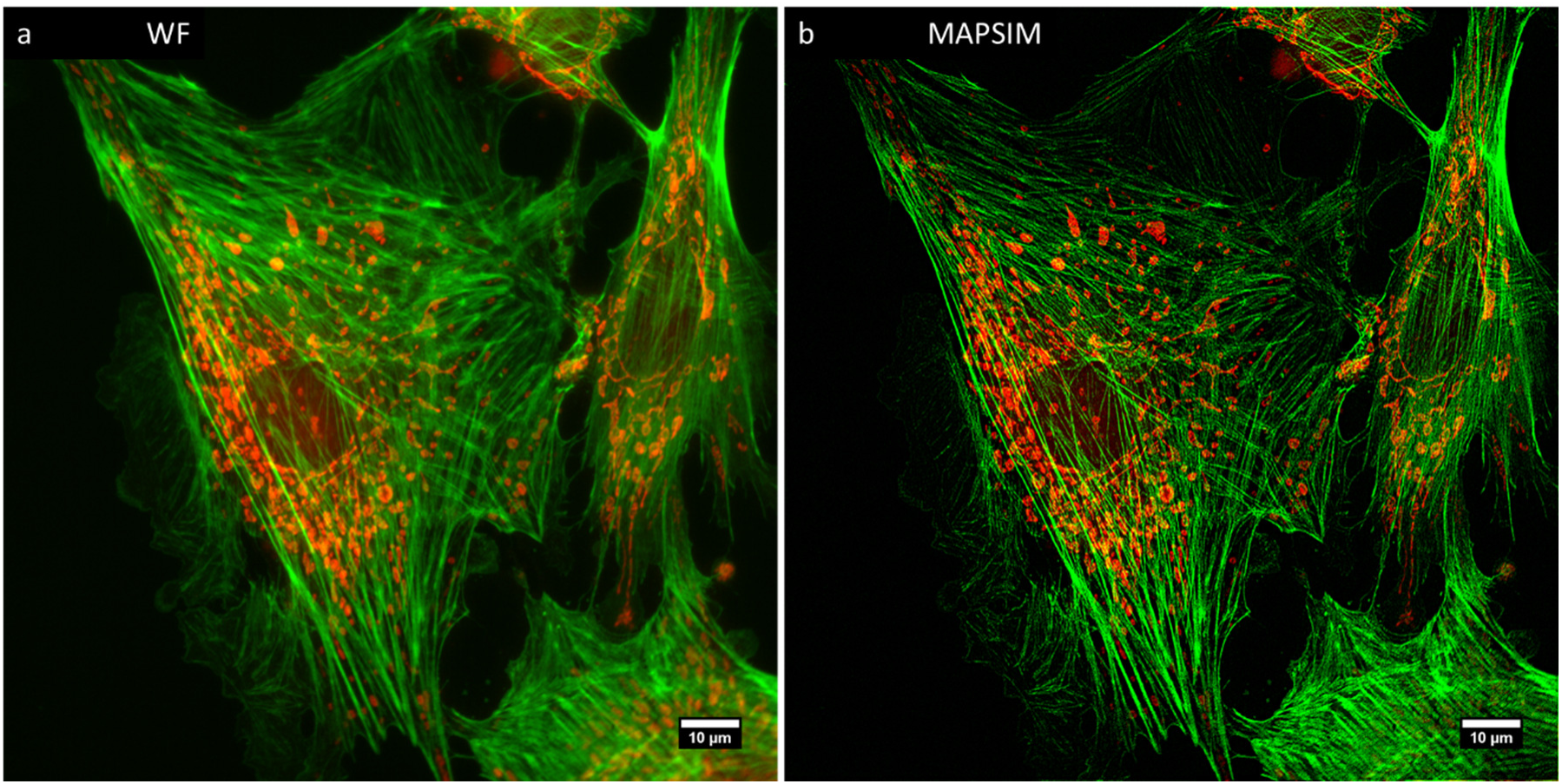
2D SIM imaging of fixed BPAE cells labeled with Alexa 488-phalloidin (actin) and mitotracker CMXRos (mitochondria). (a) WF, (b) MAP-SIM.

## 5. Discussion

SIM results sometimes suffer from artifacts related to the illumination pattern. The artifacts, which can be severe and are a cause for concern, can be due to several factors including illumination pattern phase instability and pattern distortion because of refractive index mismatch between the sample and the immersion fluid. In our hands, MAP-SIM results do not suffer from detectable patterned artifacts, Fig. 4(c), and the FFT of the MAP-SIM result is free of noticeable spurious peaks, Fig. 4(f). We attribute this to several factors, primarily the use of incoherent illumination together with a SLM for pattern generation. This, combined with precise synchronization of the SIM system helps eliminate patterned artifacts. Additional artifacts in SIM images can arise due to the detector. In sCMOS cameras like the one we used, each pixel reads out through its own amplifier and as such, each pixel exhibits a different gain. While very minor, such artifacts can be corrected using a variance stabilization method as has been introduced for single molecule localization microscopy [53].

There are several other advantages to the use incoherent illumination in SIM, including removing the need for a pupil plane mask to block unwanted diffraction orders. Also, incoherent imaging of a microdisplay for pattern formation means that the pattern spatial frequency in the sample plane does not depend on the wavelength of the light which is used. On the other hand, in incoherent illumination SIM such as we used here, the contrast of the illumination pattern decreases with increasing spatial frequency according to the incoherent optical transfer function [53]. In coherent illumination SIM [8,9,11], the coherent optical transfer function applies [53], and so the pattern contrast does not decrease with increasing spatial frequency. This means that coherent illumination SIM can more efficiently mix high resolution information from outside the frequency limit into the detection passband of the microscope, thereby potentially achieving better resolution than what we achieved in this work. We achieved a lateral resolution enhancement factor of ~1.8 (Fig. 2), whereas a factor of 2.0 is expected for coherent illumination SIM.

The LCOS microdisplay (and vendor-supplied microdisplay-timing program) we used can display an illumination pattern and switch to the next pattern in the sequence in 1.14 ms, allowing unprocessed SIM images to be acquired at rates of approximately 875 Hz. However, such rapid imaging is not useful if the reconstructed SIM images are of poor quality, for example if they suffer from low signal to noise ratios. Specifying the fastest possible acquisition rate is inadequate without consideration of the resolution and SNR of the results. Our resolution analysis shown in Figs. 2-4 (see also supplementary figure 4) uses measured quantities to evaluate SIM results and helps to make realistic conclusions about imaging speeds.

## 6. Re-use potential

The presented SIM datasets can be reused in several ways. Researchers investigating SIM reconstruction algorithms can use the datasets to compare their results with those presented here, including the newer method MAP-SIM. Also, the data may be further analyzed in other ways. One possibility is shown in the supplementary material (part 2: single particle tracking experiments in LAMP1-GFP cells.) Here, we used single particle tracking methods to study the mobility of lysosomes within U2-OS cells.

## Availability of source code and requirements

Project name: SIMToolbox v1.3

Project home page: http://mmtg.fel.cvut.cz/SIMToolbox/

Operating system: platform independent

Programming language: MATLAB

License: GNU General Public License v3.0

## Detailed software compatibility notes

The SIMToolbox GUI was compiled with MATLAB 2015a and tested in Windows 7 and 8. The GUI is a stand-alone program and does not require MATLAB to be installed. To use the MATLAB functions within SIMToolbox (i.e., without the GUI), MATLAB must be installed. The functions were mainly developed with 64 bit MATLAB versions 2012b, 2014a, 2015a in Windows 7. When using SIMToolbox functions without the GUI, the MATLAB “Image Processing Toolbox” is required. SIMToolbox also requires the “MATLAB YAML” package to convert MATLAB objects to/from YAML file format. Note that this package is installed automatically when using the GUI.

## Availability of data

All raw and analyzed data is available on GigaDB at http://gigadb.org/site/index.

## Abbreviations

GFP: green fluorescent protein
NA: numerical aperture
PSF: point spread function
WF: wide field
SIM: structured illumination microscopy
PSD: power spectral density
PSD_ca_: circularly averaged power spectral density
SNR: signal to noise ratio
SBR: signal to background ratio

## Ethics approval and consent to participate

Not applicable

## Consent for publication

Not applicable

## Author Contributions

TL: analyzed data, developed computer code; JP: analyzed data, developed computer code KF: supervised research; KS: analyzed data JB: acquired data; GH: conceived project, acquired data, analyzed data, supervised research, wrote the paper

## Acknowledgements

The authors thank Dr. Donna Arndt-Jovin and Dr. Tomas Jovin of the Max Planck Institute for Biophysical Chemistry (Göttingen, Germany) for the A431 cells. The authors thank Viola Hausnerová and Christian Lanctôt, Charles University in Prague (Prague, Czech Republic), for the LAMP1-GFP cells. The authors thank Pavel Křížek, Zdeněk Švindrych, and Martin Ovesný for help with data acquisition, microscopy development, programming, and data analysis.

## Supplementary Information

### 1. OS-SIM and MAP-SIM

#### 1.1 Optical Sectioning SIM (OS-SIM)

Several data processing methods are possible for generating optically sectioned images from SIM data (OS-SIM) [1,2]. The most familiar implementation of this technique was introduced in 1997 by Neil et al. [3]. Their method works by projecting a line illumination pattern onto a sample, followed by acquisition of a set of three widefield images with the pattern shifted by relative spatial phases 0, 2π/3, and 4π/3. An optically sectioned image can be recovered computationally as

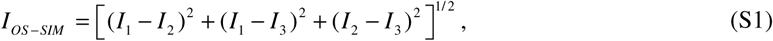

where *I*_*OS*-*SIM*_ is optically sectioned image, and *I_1_*, *I_2_*, and *I_3_* are the three images acquired with different pattern positions. This is sometimes called a ‘square law’ method. If the sum of the individual SIM patterns results in homogeneous illumination, as is the case in our setups, a widefield image can be recovered from SIM data as an average of all images:

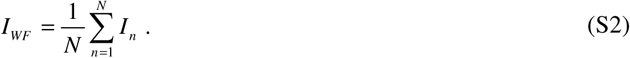

This is the approach we used throughout this study to generate WF images.

In this study we used a method known as homodyne detection [1–3].

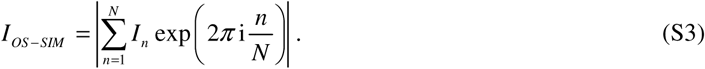

We previously showed that this processing method offers results with better optical sectioning than the square law method of Eq. S2 [2].

#### 1.2 SIM with maximum a posteriori probability estimation

MAP-SIM has been described previously [4]. In this case, image acquisition in structured illumination microscopy can be described as

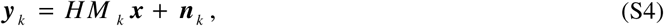

where *M_k_* represents the *k*-th illumination pattern, ***y**_k_* denotes a low-resolution (LR) image acquired using the *k*-th illumination pattern, ***x*** is an unknown, high-resolution (HR) image, and ***n**_k_* is additive noise. *H* is a matrix describing the convolution between the HR image and the point spread function (PSF) of the system. The position of the illumination patterns in the camera images were determined using a calibrated camera according to our previous work [2]. We model the PSF as an Airy disk, which in Fourier space leads to an optical transfer function (OTF) of the form [5]

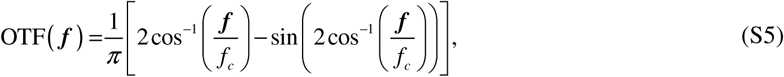

where ***f*** is the spatial frequency. We estimate the cut-off frequency *f_c_* by calculating the radial average of the power spectral density (PSD) of a widefield image of 100 nm fluorescent beads.

Using a Bayesian approach [4,6–10], high-resolution image estimation can be expressed as a minimization of a cost function according to

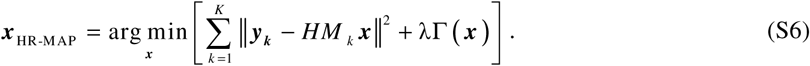

The cost function in Eq. S6 consists of two terms. The first term describes the mean square error between the estimated HR image and the observed LR images. The second term is a regularization term. To ensure positivity and promote a smoothness condition on the HR image, we rely on quadratic regularization composed of finite difference approximations of the first order derivative at each pixel location [11]

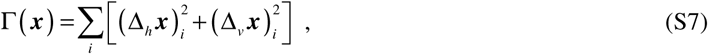

where Δ_*h*_ and Δ_*v*_ are the finite difference operators along the horizontal and vertical direction of an image, and (·)_*i*_ denotes the *i*-th element of a vector. The contribution of **Γ**(***x***) is controlled by the parameter λ, a small positive constant defining the strength of the regularization (typically λ = 0.01). We solve Eq. (S6) using gradient descent methods.

#### 1.3 Spectral merging

MAP estimation of high resolution images obtained with structured illumination enables reconstruction of high resolution images (HR-MAP) with details unresolvable in a widefield microscope. However, MAP estimation as described above does not suppress out of focus light. On the other hand, the homodyne detection method

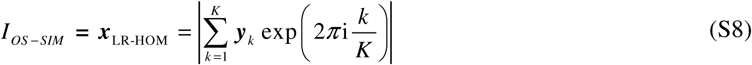

used in optical sectioning SIM (OS-SIM) [2,3] provides images (LR-HOM) with optical sectioning but with only a slight improvement in lateral resolution. Noting that the unwanted out of focus light is dominant at low spatial frequencies, we merge the LR-HOM and HR-MAP images in the frequency domain to obtain the final HR image (MAP-SIM). Frequency domain Gaussian low pass filtering is applied to the LR-HOM image and a complementary high pass filter is applied to the HR-MAP image. We use a weighting scheme which can be described by

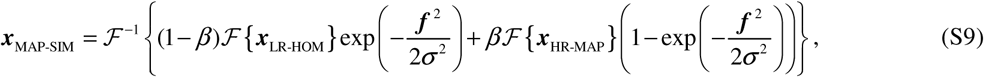

where 𝓕, 𝓕^−1^ denotes the Fourier transform operator and its inverse, respectively, ***f*** is the spatial frequency, ***σ*** is the standard deviation of the Gaussian filter, and *β* is a weighting coefficient. Usually we set *β* = 0.85. We typically use a circularly symmetric cosine bell apodizing function to shape the final MAP-SIM spectrum before the final inverse FFT. The spectral merging method is shown schematically in figure S2.

**Fig. S1.**
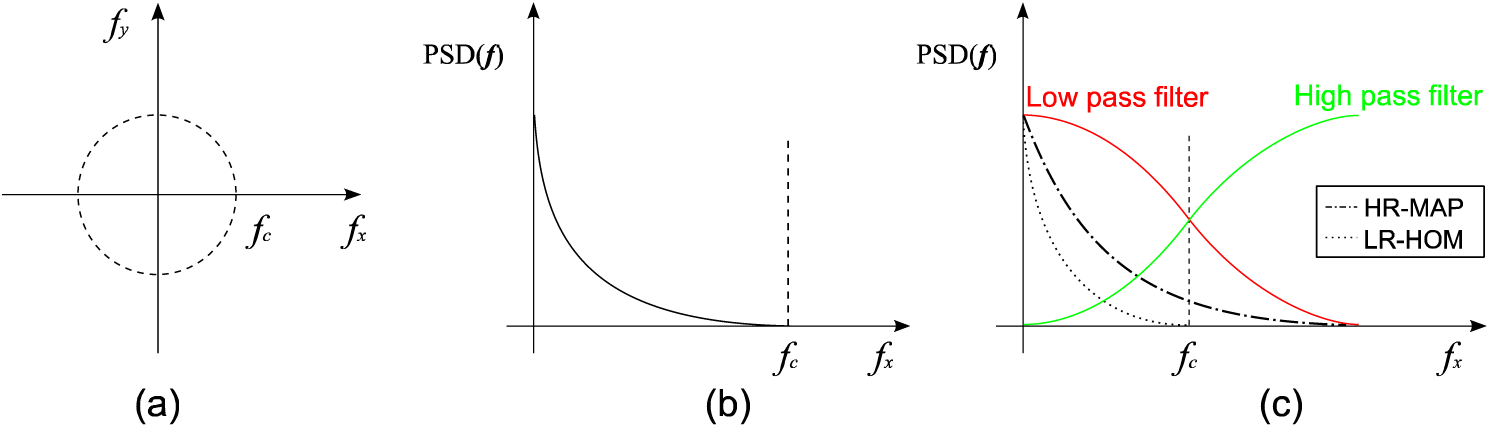
Schematic of spectral merging (a) Spatial frequencies in Fourier space, where *fc* is the cut off frequency for a WF microscope. (b) Power spectral density (PSD) in relation to the spatial frequency. (c) Blending frequency spectra of HR-MAP estimation and LR homodyne detection using low and high pass filters.

### 2. Example of data re-use: Single particle tracking experiments in LAMP1-GFP cells

#### 2.1 The optical sectioning effect of SIM allows tracking of low signal to background ratio particles in LAMP1-GFP cells

Single particle tracking (SPT) is a computer enhanced microscopy method used to track the motion of biological molecules or vesicles [12–14]. In SPT, a particle trajectory is obtained from position coordinates over a series of time steps. There are three basic steps in single particle tracking analysis [15]. The first is detection of the particles in the raw data. This may be regarded as segmentation or feature detection. The second step is localization of the particle, usually accomplished by fitting a small region of interest (7×7 pixels in our case) to a two-dimensional Gaussian function. The third step is to link the localizations together from one frame to the next to create a particle trajectory which is as long as possible. These three steps combine to determine whether a particle can be successfully tracked. If a particle is not detected in every frame through the sequence, the trajectory will be truncated at the point where the particle was lost. MAP-SIM offers very high optical sectioning ability [4]. Because of this, the signal to background ratio (SBR) of the particles is higher, and the particles are thereby much easier to detect in the images. Our single particle tracking algorithm [16] is state of the art, however, in this particular case, it is unable to detect dim, faster moving particles in the widefield data consistently enough to be able to build a trajectory longer than a few frames. However, using MAP-SIM, we were able to successfully track these particles.

In this experiment, trajectories of single LAMP1-GFP particles (lysosomes, endosomes, or other vesicles containing LAMP1-GFP), were obtained by a SPT algorithm implemented in MATLAB [16]. Briefly, the intensity average of the reconstructed WF or MAP-SIM image stack was subtracted from each individual image within the stack to reduce sCMOS camera-induced fixed pattern noise and for feature enhancement. This is shown in Fig. S2(a). Single particle trajectories were determined from the processed data sets by selecting the initial starting particle coordinates by hand, Gaussian fitting of the imaged particle, and building trajectories from coordinates based on determining the probability of finding a diffusing particle in two dimensions at a given distance from its starting point after a given time [16]. The particles were tracked for at least 12 up to a maximum of 132 time steps of 250 ms each. Fig. S2(b) shows an example trajectory of a tracked particle. After this process of building uninterrupted trajectories, the mean-squared displacement (MSD), was calculated. The MSD is a measure of the average speed a particle travels and is calculated for each time difference Δ*t* in the track. The MSD plot was computed up to *n*Δ*t* < 1/4 of the total number of acquired time frames, where *n* is the number of available displacements of a given duration *n*Δ*t* in the track record [12,16].

We tracked 60 LAMP1-GFP particles in the MAP-SIM image sequence taken from Fig. 4 of the main paper. Figure S3(a) depicts the corresponding MSD plots and Fig. S3(b) shows a histogram of the number of particles with a given hop speed computed from the average distance traveled during the shortest lag time, Δ*t* = 250 ms.

We attempted to track the same particles in the WF image sequence using the same initial particle coordinates and tracking algorithm. We kept all algorithm settings the same between the two data sets except for the size of the point spread function. Of the 60 particles tracked in MAP-SIM, 39 or 65% were successfully tracked in the WF data. Figure S3(d) depicts the corresponding MSD plots obtained from the WF single particle trajectories and Fig. S3(e) shows a histogram of the number of particles with a given hop speed. Comparing the MAP-SIM MSD plot, Fig. S3(a), to the WF MSD plot, Fig. S3(d), it is evident that fast moving LAMP1-GFP particles, represented by steeper MSD curves, cannot be tracked as successfully in the WF data. As noted above, MAP-SIM increased the SBR of the particles we attempted to track. This is shown in Figs. S3(c) and S3(f). To evaluate the SBR of the particles, we evaluated a region of interest (ROI) around each particle (9×9 pixels). We calculated the local SBR as the ratio between the maximum (average of the 5% of the highest pixel values) and minimum (average of the 5% of the lowest pixel values) in the region. To analyze single particle tracking (SPT) experiments, we used custom routines [16,17] in MATLAB using DIPimage [18].

**Fig. S2.**
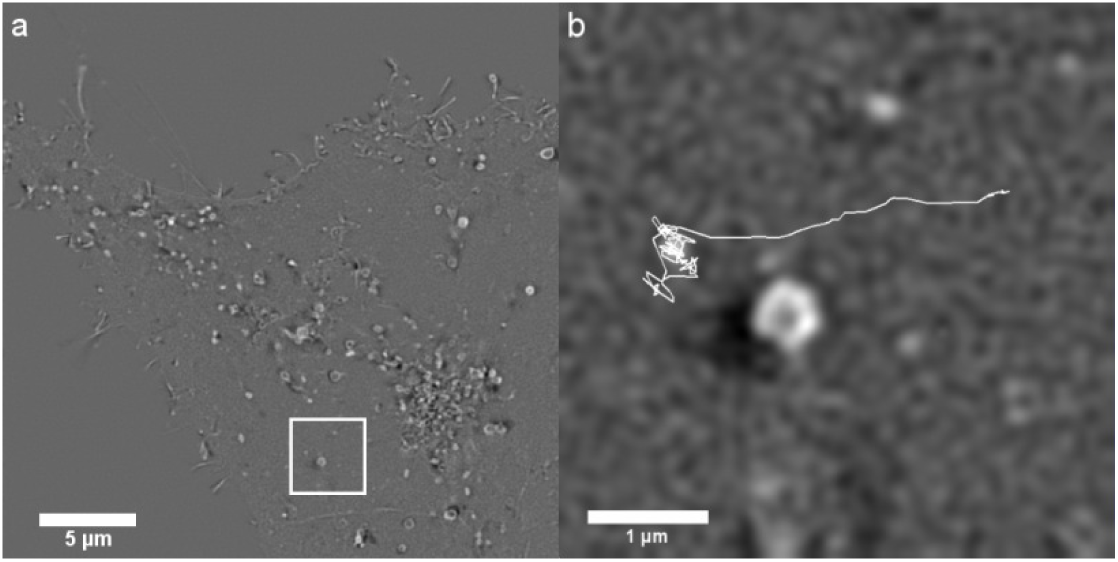
Single particle tracking. (a) MAP-SIM image from Fig. 2(c) after subtraction of the average of the stack of images. (b) LAMP1-GFP particle trajectory (132 frames) within the boxed region in (a). The particle exhibits confined diffusion followed by directed movement.

**Fig. S3.**
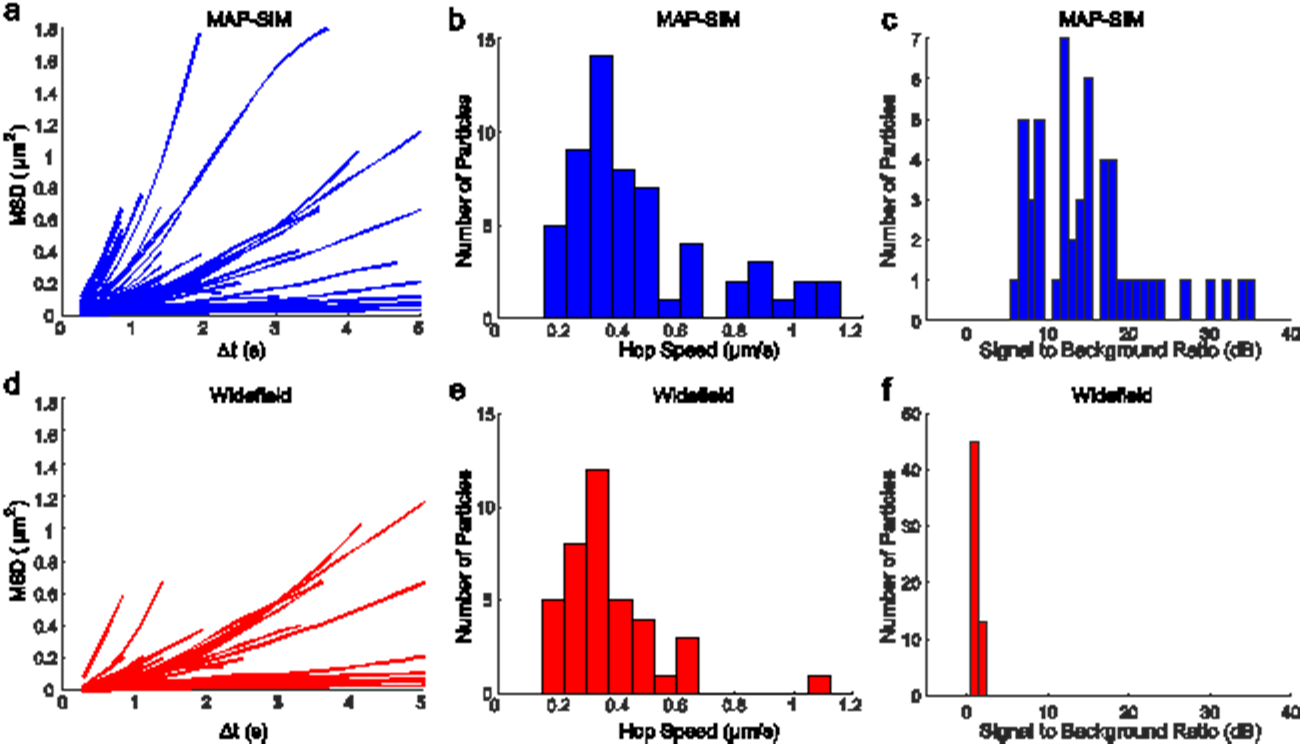
MSD plot of (a) 60 LAMP1-GFP particle trajectories obtained from a MAP-SIM image sequence and (d) 39 LAMP1-GFP particle trajectories obtained from WF image sequence. (b) and (e) histograms of particle hop speeds computed for the average distance traveled during the time lag Δt = 250 ms. (c) and (f) histograms of the signal to background ratio calculated for each particle that was tracked in the MAP-SIM (c) and WF (f) data.

### 3. Changing the exposure time affects the resolution achieved in SIM experiments

As noted in the main paper, the FLCOS microdisplay (and vendor-supplied microdisplay-timing program) we used can display an illumination pattern and switch to the next pattern in the sequence in 1.14 ms, allowing unprocessed SIM images to be acquired at rates of approximately 875 Hz. An exposure time of about 1 ms would be needed to achieve this imaging rate. However, such rapid imaging is not useful if the reconstructed SIM images are of poor quality, for example if they suffer from low signal to noise ratios (SNR). Specifying the fastest possible acquisition rate is thus inadequate without consideration of the resolution and SNR of the results.

Figure S4 demonstrates the effect of varying the camera exposure time on PSD_ca_ and thus on the spatial resolution. We imaged LAMP1-GFP cells and varied the camera exposure time from 10 ms to 100 ms per SIM sub-image. Low signal to noise ratios (in this case due to a short exposure time) causes a loss of fine details and therefore a reduction in effective resolution as estimated by our Fourier domain method [17]. These findings are not surprising, but the relationship between SNR (and therefore exposure times and usable imaging rates) and image resolution is typically not discussed in the SIM literature. This effect sometimes leads to very high noise and lower effective resolution when trying to push SIM imaging rates.

**Fig. S4.**
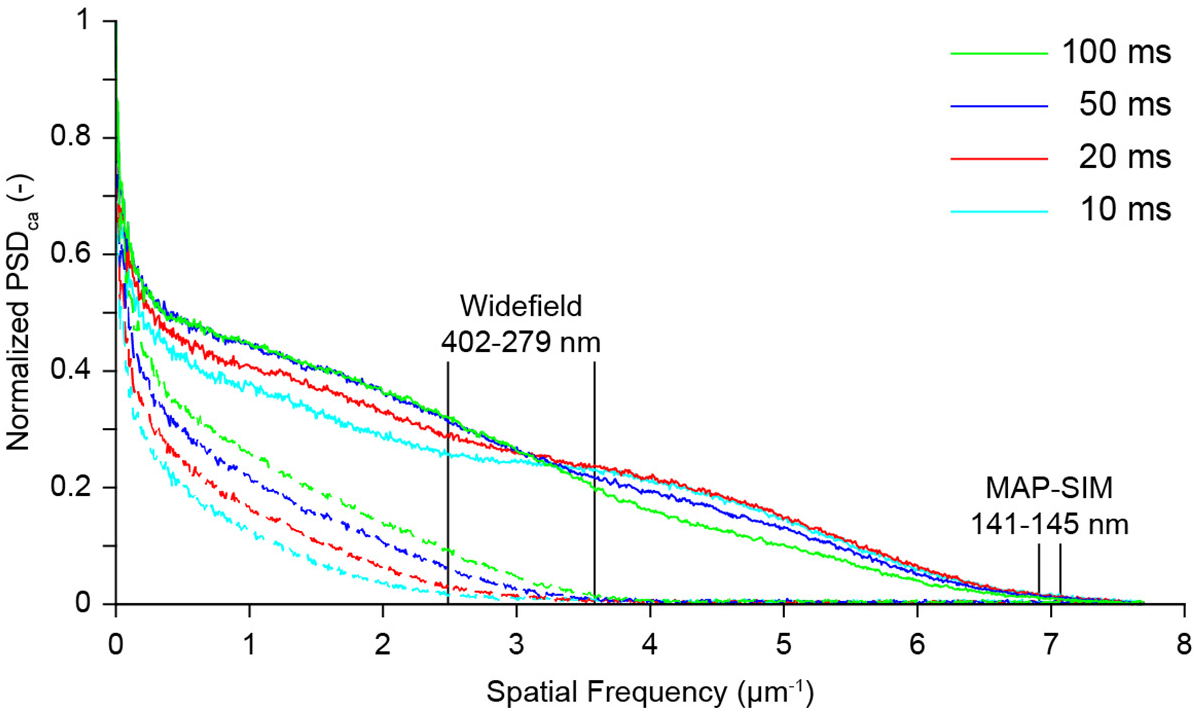
Normalized circular average power spectral density (PSD_ca_) measured as a function of exposure time for WF and MAP-SIM.

### 4. Comparison of MAP-SIM and spinning disk confocal microscopy

We also used LAMP1-GFP cells to compare MAP-SIM with a spinning disk confocal microscope (Andor Revolution system with Olympus IX81 microscope and UPLSAPO 100×/1.40 NA oil immersion objective). The results indicate a resolution of 342 nm for spinning disk confocal, 279 nm for WF, 286 nm for OS-SIM, and 145 nm for MAP-SIM.

**Fig S5.**
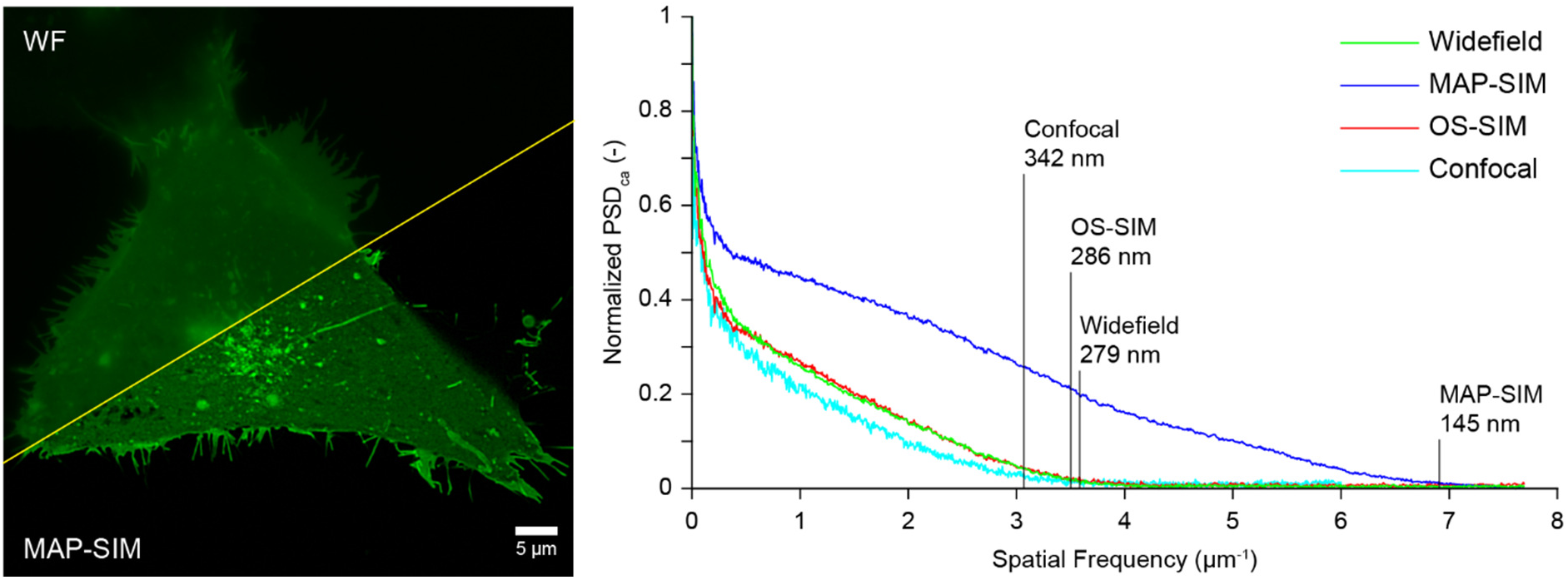
(at left) U2-OS cells expressing LAMP1-GFP, WF and MAP-SIM. (at right) PSD measurements for WF, OS-SIM, MAP-SIM, and spinning disk confocal.

